# Viral Architecture of SARS-CoV-2 with Post-Fusion Spike Revealed by Cryo-EM

**DOI:** 10.1101/2020.03.02.972927

**Authors:** Chuang Liu, Yang Yang, Yuanzhu Gao, Chenguang Shen, Bin Ju, Congcong Liu, Xian Tang, Jinli Wei, Xiaomin Ma, Weilong Liu, Shuman Xu, Yingxia Liu, Jing Yuan, Jing Wu, Zheng Liu, Zheng Zhang, Peiyi Wang, Lei Liu

## Abstract

Since December 2019, the outbreak of Coronavirus Disease 2019 (COVID-19) spread from Wuhan, China to the world, it has caused more than 87,000 diagnosed cases and more than 3,000 deaths globally. To fight against COVID-19, we carried out research for the near native SARS-CoV-2 and report here our preliminary results obtained. The pathogen of the COVID-19, the native SARS-CoV-2, was isolated, amplified and purified in a BSL-3 laboratory. The whole viral architecture of SARS-CoV-2 was examined by transmission electron microscopy (both negative staining and cryo-EM). We observed that the virion particles are roughly spherical or moderately pleiomorphic. Spikes have nail-like shape towards outside with a long body embedded in the envelope. The morphology of virion observed in our result indicates that the S protein of SARS-CoV-2 is in post-fusion state, with S1 disassociated. This state revealed by cryo-EM first time could provide an important information for the identification and relevant clinical research of this new coronavirus.

## Introduction

Coronaviruses (CoV) are a large family of zoonotic viruses that could be transmitted from animals to humans and cause illness ranging from common cold to severe diseases [1], for instance, Middle East Respiratory Syndrome (MERS) [2] [3] and Severe Acute Respiratory Syndrome (SARS) [4] in human being. Since the early December 2019, the outbreak of an unknown epidemic pneumonia in Wuhan, a city in Hubei Province of China, is identified to be caused by a novel coronavirus [5], named as SARS-CoV-2 by the International Committee on Taxonomy of Viruses (ICTV) and the disease was named as Coronavirus Disease 2019 (COVID-19) by The World Health Organization (WHO) [6]. Up to now, nearly 80,000 confirmed cases and more than 2,800 deaths in China have been reported (from WHO website). Rapid increasing number of cases also has been reported by many other countries around the world. The globe is now facing a big threat of COVID-19 [7]. While the world are jointing efforts to develop diagnostics, therapeutics, and vaccines to fight against COVID-19, information about cultivation, purification and super molecular structure of SARS-CoV-2 in their native state is in need urgently.

CoV are enveloped single-stranded positive-RNA viruses. General knowledges of the super molecular structure of the virions have been advanced by determining several kinds of CoV using electron microscope and X-ray crystallography in the past decades. Coronaviruses have a spherical or moderately pleiomorphic virions. The virion’s diameter ranges from 80 nm to 120 nm. Nucleic acid and nucleocapsid protein of coronaviruses are tightly packed in virion. Surface of virion is lipid bilayer hijacked from host cell, on which are located several important structure proteins, including spike (S) protein, envelope (E) protein and membrane (M) protein [8]. S protein, including S1 subunit and S2 subunit, is considered as the most important protein in CoV. The S1 subunit facilitates attachment to the host cells and the S2 subunit is involved in subsequent fusion of the virus and host membrane [9]. Several recent studies considering the structure of SARS-CoV-2 were all focused on the S protein. *Wrapp et al.* [10] reported a structure at 3.5 Å resolution of SARS-CoV-2 S protein. *Yan et al.* [11] reported the complex structure of SARS-CoV-2 S protein with human host cell binding receptor angiotensin-converting enzyme 2 (ACE2). *Lan et al.* [12] reported crystal structure of SARS-CoV-2 S protein’s receptor binding domain (RBD) region binding with ACE2. However, all proteins mentioned above are engineered in laboratory by recombinant expression system, not from real virus and the structure as whole of virion is thus still lacking.

Furthermore, safety is one of the major factors that restricts virus study of SARS-CoV-2 due to a high risk of possible transmission, live viruses must be operated in Biosafety Level 3 (BSL-3) laboratory or above. Here we report the successful isolation and purification of SARS-CoV-2 in BSL-3 laboratory and revealed the whole viral architecture of SARS-CoV-2 by transmission electron microscopy (both negative staining and cryo-EM). It is the first time to image the near native SARS-CoV-2, the pathogen of COVID-19, by cryo-EM.

## Results and Discussion

A 62-year-old male was admitted to our hospital on 15 January, 2020 with pneumonia, and was further diagnosed as COVID-19. An epidemiological investigation also confirmed a Wuhan travel history between 1 January and 14 January of this patient, and the symptoms started on 11 January, 2020, including fever and cough. Common respiratory viruses including influenza A virus, influenza B virus, adenoviruses, human parainfluenza virus and other human coronaviruses were tested to be negative. Lymphopenia, elevated CRP and IL-6 were found upon admission (Table S1). CT scans showed multiple ground-glass opacities and in bilateral lungs at the early stage, and lung consolidation occurred during hospitalization (Figure S1).

The bronchoalveolar lavage fluid (BALF) sample was collected and subjected to next-generation sequencing and virus isolation was performed in the BSL-3 (Biosafety level 3) laboratory. Typical cytopathic effects (CPE) were observed at 4 days post inoculation (dpi) in Vero cell, including cell rounding, shrinkage, lysis, and detachment throughout the cell monolayers (Figure 1A). Viral RNAs could be detected in the cell culture supernatant using a CFDA approved commercial kit with low Ct values (Figure 1B). The purified SARS-CoV-2 particles showed specific reaction activity to convalescent plasmas from SARS-CoV-2-infected patients (Figure 1C) and the specific human monoclonal antibodies against the RBD of the S protein using ELISA assay (Figure 1D). Meanwhile, the virus could also be detected by immunofluorescence using the patient’s plasma (Figure 1E). The genome sequence of this virus has been submitted to the Global Initiative on Sharing Avian Flu Data (GISAID) with an access number of EPI_ISL_406594, and designated as “BetaCoV/Shenzhen/SZTH-003/2020”. Phylogenetic analyses showed that the viruses possessed high homology with the other isolates, and two closest isolates are BetaCoV/Wuhan/IPBCAMS-WH-04/2019 from Wuhan and SARS-CoV-2/NTU01/2020/TWN from Taiwan (Figure 1F).

**Figure 1.**
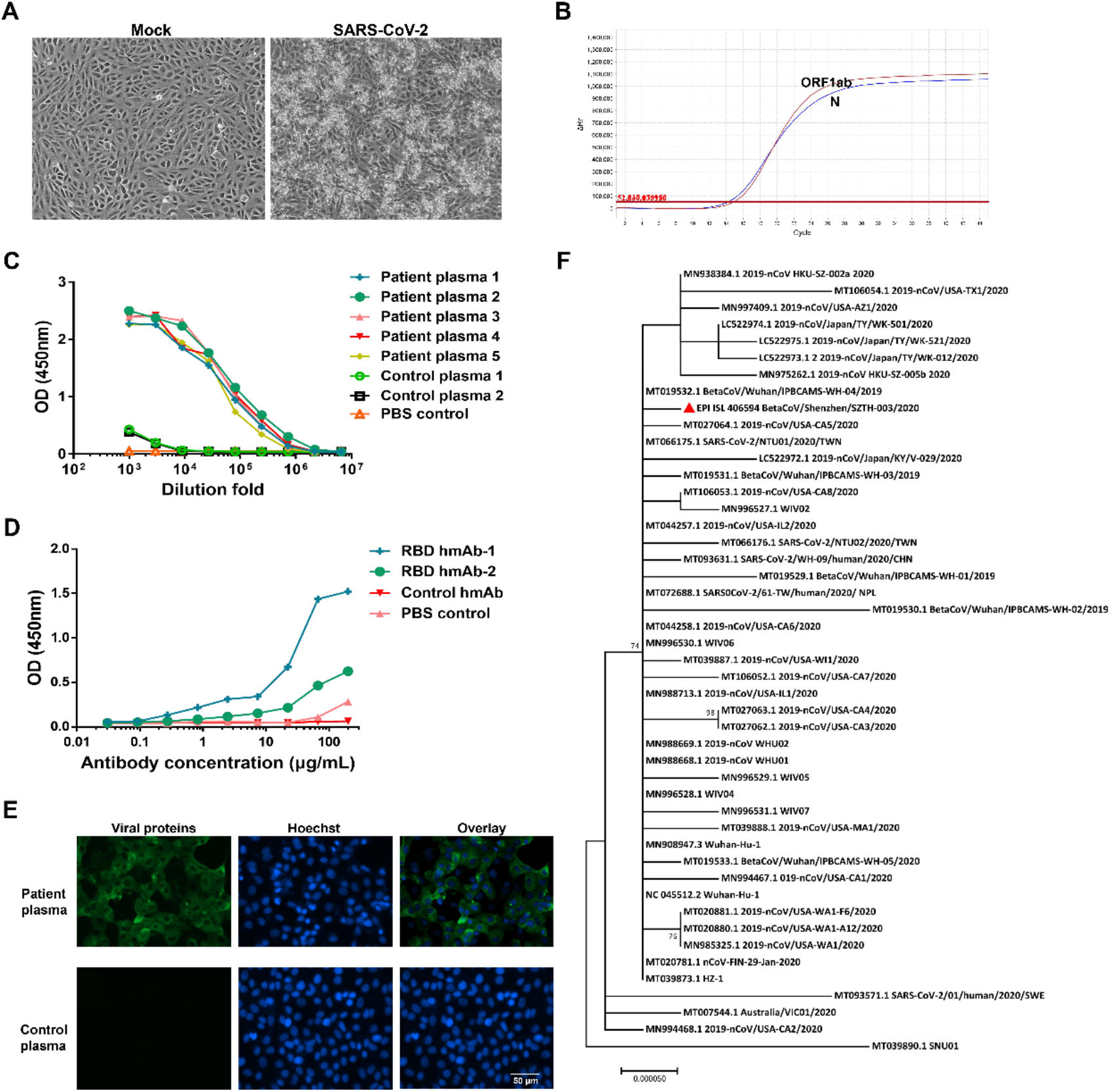
Isolation and identification of the SARS-CoV-2 virus. (A) Vero cell was inoculated with BALF sample, and the CPEs were observed at 4-dpi. (B) RNAs were extracted from the cell culture supernatant, and detected using a commercial kit targeting the ORF 1ab (red) and N (blue) genes of SARS-CoV-2. Testing the convalescent plasma IgG antibody (C) and SARA-CoV-2-RBD specific human monoclonal antibodies (D) using the purified SARS-CoV-2 particles. The control plasmas 1 and 2 were obtained from a patient recovered from influenza A virus infection and a healthy volunteer, respectively. The control monoclonal antibody is a human monoclonal antibody specific to influenza A virus generated by our laboratory. The PBS control was serum-free. (E) Viruses were detected by IFA using the patient’s plasma, and plasma from a healthy control was used as negative control. Scale bar, 100 μm. (F) Phylogenetic characteristics of BetaCoV/Shenzhen/SZTH-003/2020, and this isolate was indicated with a triangle.

In Figure 2 and Figure 3 we present the EM observations of SARS-CoV-2, both by negative staining (Figure 2A-2E) and cryo-EM (Figure 3A-3F). The virion particles are roughly spherical or moderately pleiomorphic, with diameters range from 80nm-160nm. For majority of the virions, the periphery is well defined and enveloped by lipid bilayer, which could be clearly seen in cryo-EM image. Tightly underneath the lipid bilayer is the condensed density formed by nucleic acid and nucleocapsid protein of SARS-CoV-2. Though lots of the particles lost spike during purification or deactivation, 20%-30% percent virions still have spikes around the envelope.

**Figure 2.**
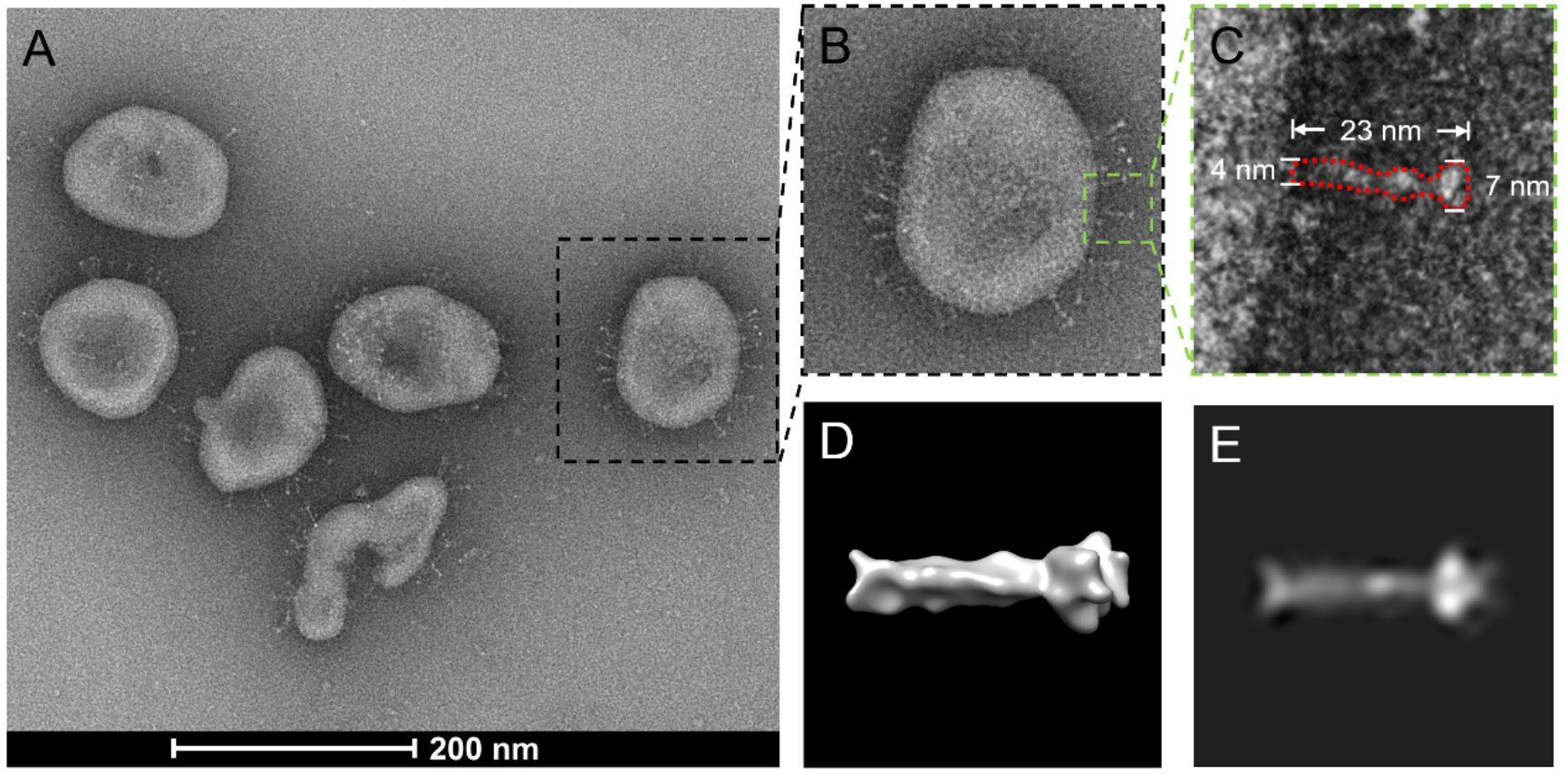
Negative stain EM results of SARS-CoV-2. (A). Image of negative stained SARS-CoV-2. Nail-like spikes can be clearly seen. (B). Enlarged view of virion boxed in (A). (C) Zoom-in view of a spike boxed in (B). The shape is depicted by red dot line. Length, the diameter of stem and spike’s head are 23nm, 4nm and 7nm, respectively. (D). Three-dimensional surface of post-fusion state S2 protein (EMDB code: 9597) [15]. (E). Projection of post-fusion state S2 protein [15].

**Figure 3.**
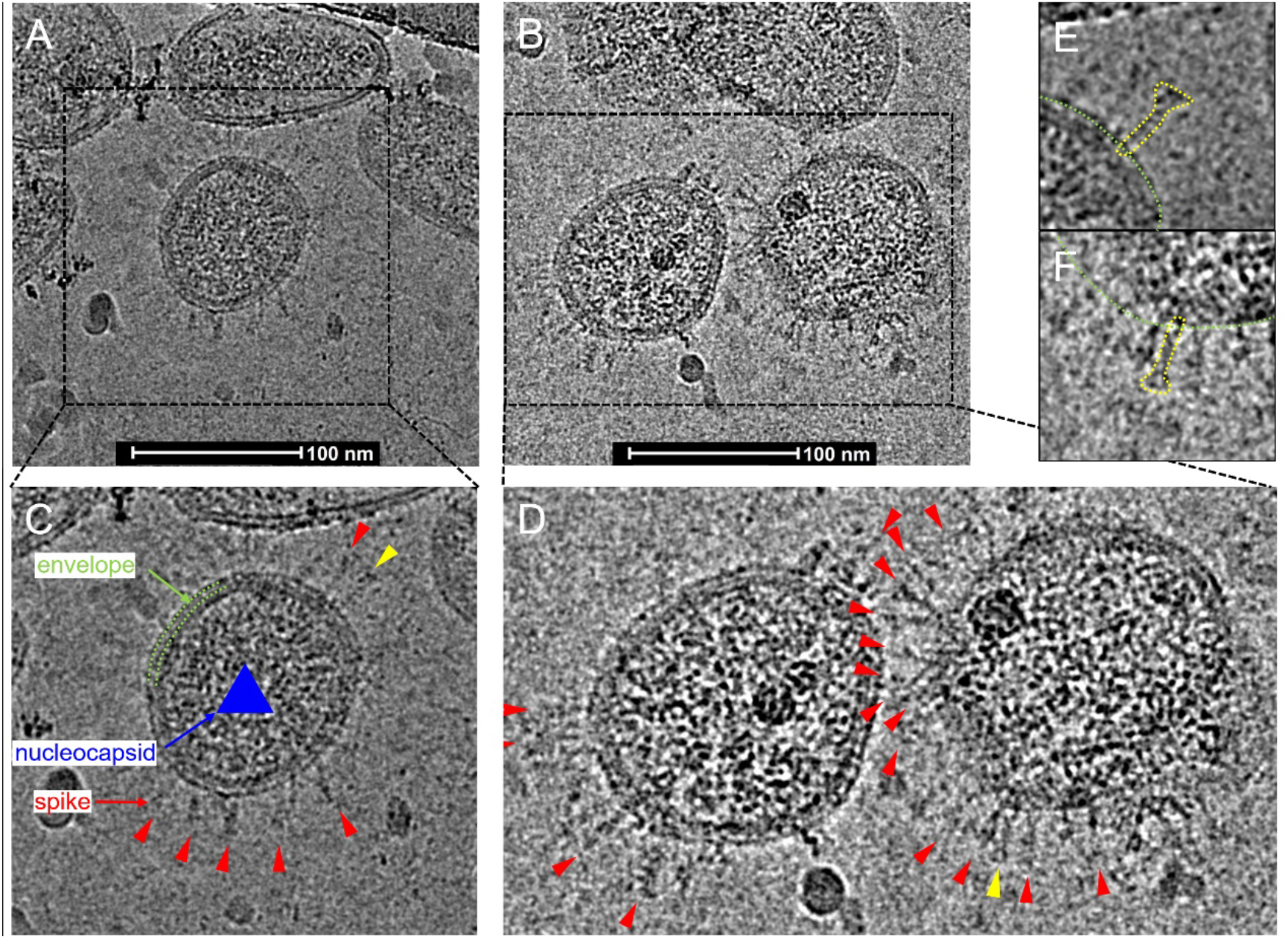
Cryo-EM results of SARS-CoV-2. (A) and (B). Cryo-EM images of SARS-CoV-2. (C). Zoom-in view of the virion showed in (A). Envelope and nucleocapsid are indicated by green and blue respectively, remarkable spikes are indicated by red triangles. (D). Zoom-in views of the two virions showed in (B), remarkable spikes are indicated by red triangles. (E). Zoom-in view of the spike indicated by yellow triangle in (C). The shape is depicted by yellow dot lines. (F). Zoom-in view of the spike indicated by yellow triangle in (D). The shape is depicted by yellow dot lines.

Different with spikes reported in previous study [5], spikes in our images are much thinner both in negative staining image (Figure 2) or cryo-EM image (Figure 3). Spikes here have nail-like shape with a long body embedded in the envelope. Width of spike’s head is around 7 nm. The length of whole spike is around 23nm. S protein of coronavirus consisted of S1 and S2 subunits. When S1 subunit is disassociated, S2 undergoes a conformation change, extending itself from compressed form to a nail-like shape, termed as post-fusion state. The nail-like shape was also observed by other researchers [13] [14] [15]. Work done by *Song et al.* [15] gave a detailed structure analysis of S2 protein using recombinant protein. Not only they reported the morphology of S2 protein at post-fusion state, they also reconstructed a 3D map. The 2D average image (Figure 2E) and 3D map (Figure 2D) reported in their paper [15], have a consistent result with our observation both in negative staining or Cryo-EM images (shown in Figure 2C and Figure 3E,3F).

## Methods

### Sample Collection and Virus Isolation

The BALF was collected from the patient at 1 day after admission. Vero, LLC-MK2 and Huh-7 cells were used for the virus isolation in the biosafety level 3 (BSL-3) laboratory. BALF sample was centrifuged at 5, 000 rpm at 4°C for 5 minutes, and then 200 μl of the supernatant was added to the monolayer of cell and incubated at 37°C and 5% CO_2_ for 1 hour. Then the cells were washed with PBS for 3 times, and the fresh DMEM containing 2% fetal bovine serum (FBS) and 1% penicillin streptomycin (PS) was added to the cell culture. Cells were maintained at 37°C and 5% CO_2_, and CPEs were monitored daily with light microscopy. Meanwhile viral RNAs were detected at 3 and 5 dpi using RT-PCR to monitor the replication of virus.

### Quantitative Reverse Transcription Polymerase Chain Reaction(qRT-PCR)

Viral RNAs from BALF and cell culture supernatant were extracted using the QIAamp RNA Viral Kit (Qiagen, Heiden, Germany). Then quantitative reverse transcription polymerase chain reaction (qRT-PCR) was performed using a commercial kit specific for 2019-nCoV detection (GeneoDX Co., Ltd., Shanghai, China) as reported previously[16].

### Indirect immunofluorescence assay (IFA)

IFA was done as described previously[17] [2]. In brief, vero cells were fixed in 4% formaldehyde at 48 hours post infection. Then cells were permeabilized in 0.5% Triton X-100, blocked in 5% BSA in PBS, and then probed with the plasma of this patient or healthy control at a dilution of 1:500 for 1 h at room temperature. The cells were washed three times with PBS and then incubated with either goat anti-human IgG conjugated with Alexa fluor 488 at a dilution of 1:500 for 1 h (Invitrogen, Carlsbad, CA). The cells were then washed and stained with hoechest-33342 (Invitrogen, Carlsbad, CA) to detect nuclei. Fluorescence images were obtained and analyzed using EVOS FL Auto Imaging System (Invitrogen, Carlsbad, CA).

### Enzyme Linked Immunosorbent Assay (ELISA)

Microtiter plates (Sangon Biotech) were coated overnight at 4°C with 2.5 × 10^3^ TCID_50_/well/100 μl purified and inactivated SARS-CoV-2 particles. The plates were washed twice with PBS containing 0.1% v/v Tween-20 (PBST) and blocked with blocking solution (PBS containing 2% w/v non-fat dry milk) for 2 hours at 37°C. The plates were then washed with PBST. The plasma were diluted to 1000-fold and human monoclonal antibody specific to SARS-CoV-2-RBD generated by our laboratory were diluted to 200 μg/ml into PBS as initial concentration, and serial 3-fold dilutions of sera was added to the wells and incubated at 37°C for 60 minutes. After three washes, 100 μl of horseradish peroxidase (HRP)-conjugated goat anti-human IgG antibody solution (Sangon Biotech) was added to each well and incubated at 37°C for 60 minutes. After washing, 100 μl of tetramethylbenzidine (TMB) substrate (Sangon Biotech) was added at room temperature in the dark. After 15 minutes, the reaction was stopped with a 2M H2SO4 solution. The absorbance was measured at 450 nm. All samples were run in triplicate. The ELISA titers were determined by endpoint dilution.

### Virus Preparation

Virus was amplified in Vero cell in an incubator at 37°C and 5% CO_2_ for 96 hours and the viral load was determined by qPCR assay. The harvested 400ml virus supernatant with a virus titer of about 1.08 × 10^5^ TCID_50_/ml (genome equivalent) was firstly inactivated by adding 0.05% (v/v) β-propiolactone and incubating at 4°C for 36 hours. Then the inactivated viruses were pelleted by centrifugation at 85,000 g, 4°C using a SW28 rotor. The pellet was dissolved at 4°C overnight in buffered saline (0.15M NaCl, 20mM HEPES, pH 6.8). The 3 ml of the suspended virus was layered on top of linear gradient of 40% (w/w) potassium tartrate, 20mM HEPES, pH 7.4 (bottom) to 15% (w/w) glycerol, 20mM HEPES, pH 7.4 (top) and subjected to isopycnic centrifugation in a SW40 rotor at 85,000 g, 4°C for 3 hours. The resulting milky band of virus was diluted with buffered saline and pelleted by centrifugation using an SW40 rotor at 85,000, 4°C for another 3 hours. Finally, the pellet was allowed to dissolve at 4°C overnight in 50 μl of buffered saline.

### Viral Genome Sequence and Phylogenetic Analyses

The extracted viral RNAs were subjected for next generation sequencing using the Illumina sequencing platform. The assembled sequence was further analyzed together with other genomes of SARS-CoV-2 downloaded from National Center for Biotechnology Information (NCBI) database. The phylogenetic analyses were done as previously reported [18].

### Negative stain EM imaging

The amount of 3μl purified samples were applied to glow-discharged grid with a continuous carbon layer (Electron microscopy China, China) for 1min. The grid was blotted by filter paper to absorb the excrescent sample. The grid was then stained with 3%(w/w) phosphotungstic acid for 1min. Redundant liquid was absorbed using filter paper. The grid was transferred to a Talos 120C transmission electron microscope (Thermo Fisher Scientific) performed at the 120 kV in low dose mode and imaged with a Ceta 16M CMOS detector (Thermo Fisher Scientific). Data were collected at a nominal magnification of 45000 ×, corresponding to a pixel size of 3.5 Å/pixel.

### Cryo-EM imaging

For cryo-EM sample preparation, 3μl of the sample was placed on a glow-discharged Quantifoil grid (R1.2/1.3). The grid was blotted and plunged into liquid ethane cooling by liquid nitrogen to frozen sample using a Vitrobot Mark IV (Thermo Fisher Scientific). Frozen grid was transferred to a Titan Krios transmission electron microscope (Thermo Fisher Scientific) performed at the 300 kV and images were recorded by a Falcon III direct electron detector (Thermo Fisher Scientific). Image stacks with 16 frames were collected in linear mode by EPU at a nominal magnification of 59000 x, corresponding to a pixel size of 1.14 Å/pixel. Total dose was estimated at about 60 e^−^/Å^2^.

## Acknowledgments

We acknowledge the work and contribution of all the health providers from Shenzhen Third People’s Hospital. We sincerely thank both Cryo-EM centre and BSL-3 laboratory of the Second Affiliated Hospital, School of Medicine, in South University of Science and Technology to provide the facilities and technical supports. We also thank Professors Dongfeng Gu and Mingzhao Xing for their valuable advices and read the manuscript.

This study received approval from the Research Ethics Committee of Shenzhen Third People’s Hospital, China (approval number: 2020-038). The Research Ethics Committee waived the requirement informed consent before the study started because of the urgent need to collect epidemiological and clinical data. We analyzed all the data anonymously.

---

**Figure S1.**
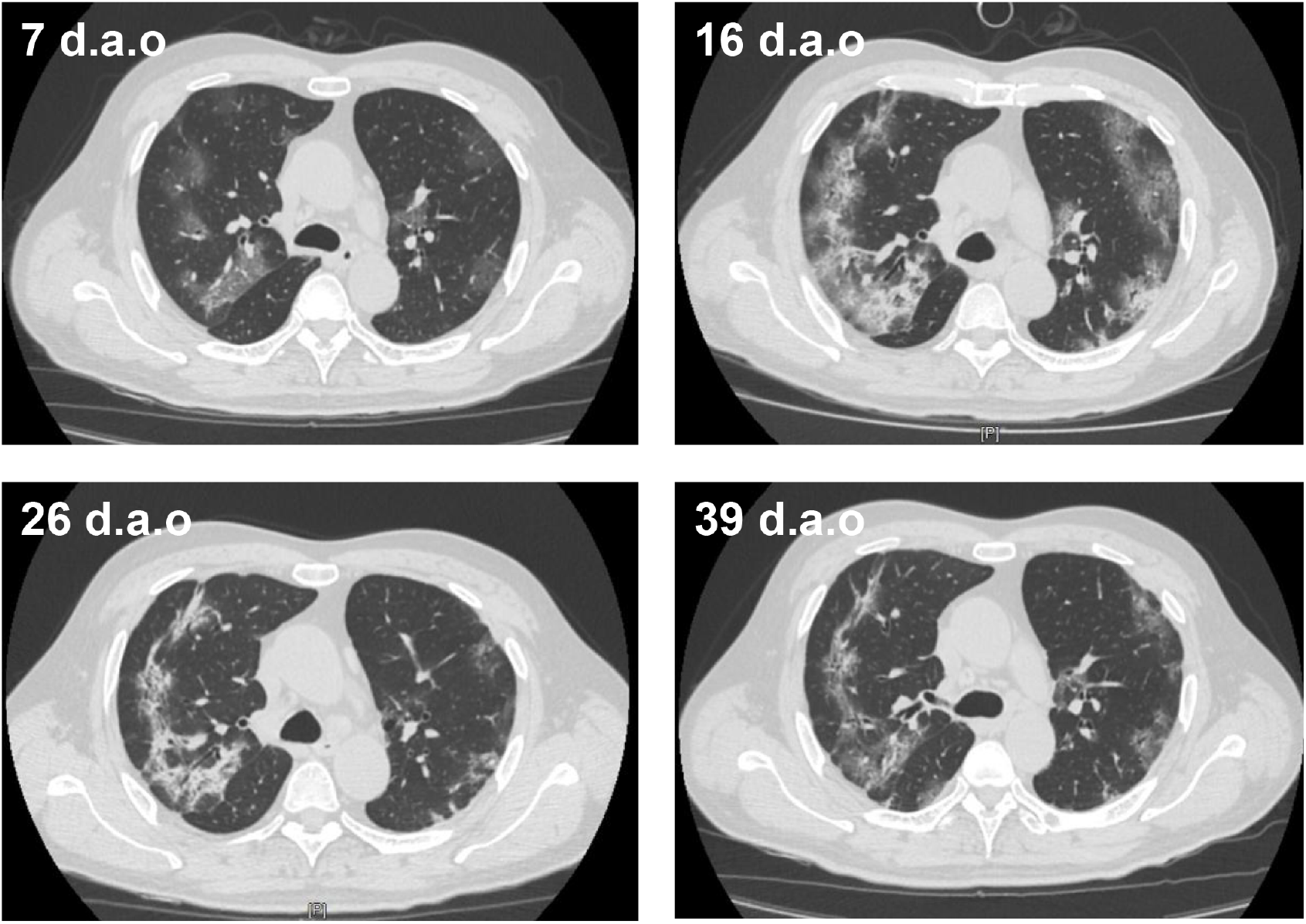
The representative CT scans of the patient at 7, 16, 26 and 39 day after illness onset (d.a.o)

**Table S1.** The clinical information of the enrolled patient.

| **Characteristics** | **Description** |
| --- | --- |
| Age | 62 |
| Gender | male |
| Disease severity | severe |
| Wuhan exposure history | Yes (1 Jan, 2020 – 14 Jan, 2020) |
| Date of symptoms onset | 11 Jan, 2020 |
| Initial symptoms | Fever/Cough |
| Date of hospitalization | 15 Jan, 2020 |
| Co-existing chronic disease | none |
| Influenza A/B viruses | - |
| Human parainfluenza virus | - |
| Respiratory syncytial virus | - |
| Human Bocavirus | - |
| Adenovirus | - |
| Human metapneumovirus | - |
| Rhinovirus | - |
| HCoV-229E/OC43/HKU1/NL63 | - |
| MERS-CoV | - |
| SARS-CoV | - |
| Interferon atomization | 16 Jan, 2020 |
| Ribavirin | 15 Jan, 2020 |
| Methylprednisolone | no |
| High-flow oxygen therapy | yes |
| Mechanical ventilation | No |
| CT finding | Bilateral pneumonia |
| White blood cell counts | $3.85 \times 10^9/\text{L}$ |
| Lymphocyte counts | $0.83 \times 10^9/\text{L}$ |
| Monocyte counts | $0.76 \times 10^9/\text{L}$ |
| D-Dimer | 0.26 $\mu\text{g}/\text{ml}$ |
| C-reaction protein | 53.0 $\mu\text{g}/\text{ml}$ |
| IL-6 | 39.5 $\mu\text{g}/\text{ml}$ |
| BALF sampling date | 21 Jan, 2020 |
| Outcome | Discharged |

